# Genome resources for climate-resilient cowpea, an essential crop for food security

**DOI:** 10.1101/059261

**Authors:** María Muñoz-Amatriaín, Hamid Mirebrahim, Pei Xu, Steve I. Wanamaker, MingCheng Luo, Hind Alhakami, Matthew Alpert, Ibrahim Atokple, Benoit Joseph Batieno, Ousmane Boukar, Serdar Bozdag, Ndiaga Cisse, Issa Drabo, Jeffrey D. Ehlers, Andrew Farmer, Christian Fatokun, Yong Q. Gu, Yi-Ning Guo, Bao-Lam Huynh, Scott A. Jackson, Francis Kusi, Cynthia T. Lawley, Mitchell R. Lucas, Yaqin Ma, Michael P. Timko, Jiajie Wu, Frank You, Philip A. Roberts, Stefano Lonardi, Timothy J. Close

**Affiliations:** Department of Botany and Plant Sciences, University of California, Riverside, USA; Department of Computer Science and Engineering, University of California, Riverside, California, USA; Institute of Vegetables, Zhejiang Academy of Agricultural Sciences, Hangzhou 310021, China; Department of Plant Sciences, University of California, Davis, California, USA; Council for Scientific and Industrial Research, Savanna Agricultural Research Institute, Tamale, Ghana; Institut de l’Environnement et de Recherches Agricoles, Saria, Burkina Faso; International Institute of Tropical Agriculture, Kano, Nigeria; Department of Mathematics, Statistics and Computer Science, Marquette University, Milwaukee, Wisconsin, USA; Institut Sénégalais de Recherches Agricoles, Thiès, Senegal; The Bill & Melinda Gates Foundation, Seattle, USA; National Center for Genome Resources, Santa Fe, New Mexico, USA; USDA-ARS Western Regional Research Center, Albany, California, USA; Department of Nematology, University of California, Riverside, California, USA; Center for Applied Genetic Technologies, University of Georgia, Athens, Georgia, USA; Illumina, Inc., San Francisco, California, USA; Department of Biology, University of Virginia, Charlottesville, Virginia, USA; Agriculture and Agri-Food Canada, Morden, Manitoba, Canada.

## Abstract

Cowpea (*Vigna unguiculata* L. Walp.) is a legume crop that is resilient to hot and drought-prone climates, and a primary source of protein in sub-Saharan Africa and other parts of the developing world. However, genome resources for cowpea have lagged behind most other major crop plants. Here we describe foundational genome resources and their application to analysis of germplasm currently in use in West African breeding programs. Resources developed from the African cultivar IT97K-499-35 include bacterial artificial chromosome (BAC) libraries and a BAC-based physical map, assembled sequences from 4,355 BACs, as well as a whole-genome shotgun (WGS) assembly. These resources and WGS sequences of an additional 36 diverse cowpea accessions supported the development of a genotyping assay for over 50,000 SNPs, which was then applied to five biparental RIL populations to produce a consensus genetic map containing 37,372 SNPs. This genetic map enabled the anchoring of 100 Mb of WGS and 420 Mb of BAC sequences, an exploration of genetic diversity along each linkage group, and clarification of macrosynteny between cowpea and common bean. The genomes of West African breeding lines and landraces have regions of marked depletion of diversity, some of which coincide with QTL that may be the result of artificial selection or environmental adaptation. The new publicly available resources and knowledge help to define goals and accelerate the breeding of improved varieties to address food security issues related to limited-input small-holder farming and climate stress.

## INTRODUCTION

Cowpea *(Vigna unguiculata* (L.) Walp.), native to Africa and a member of the Fabaceae family, is a primary source of protein in sub-Saharan Africa, where it is grown for fresh and dry grains, foliage, and forage. Cowpea is also an important crop in parts of Asia, South America, and the USA (Singh 2014). Because of its adaptability to harsh conditions, cowpea is a successful crop in arid and semiarid regions where few other crops perform well. Cowpea is important to the nutrition and income of smallholder farmers in Africa, while also contributing to sustainability of the cropping system through fixation of atmospheric nitrogen and prevention of soil erosion. Despite its relevance to agriculture in the developing world and its stress resilience, actual yields of cowpea are much lower than the known yield potential, and cowpea genome resources have lagged behind those developed for other major crop plants.

Cowpea is a diploid with a chromosome number 2n = 22 and an estimated genome size of 620 Mb (Chen et al. 2007). Its genome shares a high degree of collinearity with other warm season legumes, especially common bean *(Phaseolus vulgaris* L.) (Vasconcelos et al. 2015). Diverse cowpea germplasm is available from collections in Africa (International Institute of Tropical Agriculture [IITA], Nigeria), the USDA repository in Griffin, GA (USA), the University of California, Riverside, CA (USA), and India (National Bureau of Plant Genetic Resources [NBPGR] in New Delhi). These collections contain diversity relevant to pests, pathogens, plant architecture, seed characteristics and adaptation to marginal environments. Resources that were developed previously to support adoption of markers for breeding include a 1,536-SNP GoldenGate assay (Muchero et al. 2009), which has enabled linkage mapping and QTL analysis (e.g. Lucas et al. 2011; Muchero et al. 2013; Pottorff et al. 2014) as well as an assessment of the diversity of landraces throughout Africa (Huynh et al. 2013).

IT97K-499-35, developed at IITA, was released in Nigeria in 2008 as a line that is resistant to most races of the parasitic weed *Striga gesnerioides* that are prevalent in West Africa. This black-eyed variety has also been released as a cultivar in Mali and Ghana under the names “Djiguiya” and “Songotra”, respectively. Gene space sequences accounting for ~160 Mb of the IT97K-499-35 genome were previously published (Timko et al. 2008). In addition, 29,728 “unigene” consensus sequences, derived from 183,118 ESTs from cDNA libraries of 17 different cowpea accessions are available in the software HarvEST:Cowpea (harvest.ucr.edu) (Muchero et al. 2009).

Here we present additional resources including a physical map from BACs of IT97K-499-35, sequence assemblies for minimal tiling path (MTP) BACs and from 65x coverage whole genome shotgun (WGS) short-reads, more than 1 million SNPs discovered from sequences of 36 diverse accessions, and an Illumina Cowpea iSelect Consortium Array which represents a publicly accessible resource for screening 51,128 SNPs. These resources have been leveraged for linkage mapping and synteny analysis, used to investigate frequency and distribution of cowpea genome diversity, and used to evaluate breeding materials from four West African breeding programs, which serve one of the most food insecure regions of the world. These genomic resources do not constitute a complete sequence of the cowpea genome, yet they have enabled advances in genetic marker development, assessment of the diversity within breeding programs and comparative genomics.

## RESULTS

### Physical map and BAC sequencing

Two BAC libraries were constructed from cowpea accession IT97K-499-35 using restriction enzymes *HindIII* and *MboI* (36,864 clones each with 150 kb and 130 kb average clone insert size, respectively). High-quality BAC-end sequences (BES) were generated from 30,343 BACs using the Sanger method. BES had an average read length of 674 bp, a GC content of 37.2%, and accounted for 20.5 Mb. They were included in the WGS assembly described below. For physical mapping, 59,408 BACs (97.9% from *HindIII* and 63.2% from *MboI*) were fingerprinted using the method of Luo et al. (2003). After quality filtering, 43,717 clones were assembled into 829 contigs (40,952 BACs) and 2,765 singletons using FPC (Soderlund et al. 2000). The total number of fingerprints in the physical map represents an equivalent of 11-fold haploid genome coverage. The resulting cowpea physical map is available at http://phymap.ucdavis.edu/cowpea.

A total of 4,355 MTP clones were sequenced in combinatorial pools (Lonardi et al. 2013) using Illumina HiSeq2000. Reads were assigned to individual BACs and then assembled using SPAdes (Bankevich et al. 2012). BAC assemblies had an average N50 of 18.5 kb, an average L50 of 5.7 contigs, and a total length of 496.9 Mb (Table 1). The GC content was 34.05%. Analysis of overlap between sequenced BACs provided an estimate of non-redundant genome coverage at 372.8 Mb (~60.1% of the cowpea genome; see Experimental procedures for more details). Sequence comparison revealed that the BAC assemblies contain 17,216 (57.9%) of 29,728 cowpea EST-derived “unigene” consensus sequences available from HarvEST:Cowpea (http://harvest.ucr.edu). In addition, the BAC sequences had high homology with 15,617 (57.4%) of the 27,197 protein-coding gene models in common bean (Schmutz et al. 2014). These analyses suggest that ~40% of the cowpea genome is missing from the BAC assemblies.

**Table 1.** **BAC assembly characteristics and anchoring to the genetic map.** Values for N50 through L90 are averages of all BAC assemblies.

| SPAdes Assembly | # BACs | Total Size (bp) | Non-N Size (bp) | Avg. N50 | Avg. L50 | Avg. N90 | Avg. L90 | Avg. Longest Contig | Pv. Gene Models |
| --- | --- | --- | --- | --- | --- | --- | --- | --- | --- |
| BACs (all) | 4,355 | 496,856,182 | 496,750,728 | 18,503 | 5.74 | 4,640 | 20.39 | 30,299 | 15,617 |
| Anchored BACs | 3,502 | 419,843,633 | 419,754,909 | 20,590 | 4.13 | 5,156 | 14.51 | 33,208 | 15,061 |

N50: length for which the collection of contigs of that length or longer contains at least half of the sum of the lengths of all contigs in the BAC assembly.

N90: length for which the collection of contigs of that length or longer contains at least 90% of the sum of the lengths of all contigs in the BAC assembly.

L50: minimum number of contigs accounting for more than 50% of the BAC assembly.

L90: minimum number of contigs accounting for more than 90% of the BAC assembly.

Pv.: *Phaseolus vulgaris*.

### Whole-genome shotgun sequencing and assembly

A WGS approach using short-read sequencing was followed to assemble sequences of the cowpea genome. WGS data from IT97K-499-35 included 394 million paired-end short reads for a total of 40.6 Gb of sequence data (~65x coverage) from Illumina GAII, and Illumina HiSeq sequences from one 5 kb long-insert paired end (LIPE) library. These two datasets were assembled using SOAPdenovo (Luo et al. 2012) together with the Sanger BES described above and the “gene-space” sequences available from Timko et al. (2008). The resulting assembly has over 600,000 scaffolds (97,777 of 1 kb or longer), accounting for 323 Mb of the cowpea genome (724 Mb of total scaffold length including Ns; Table 2). This highly-fragmented assembly reflects the short length of the reads and the expected highly-repetitive genome; its close-relatives common bean (Schmutz et al. 2014) and adzuki bean (Yang et al. 2015) are ~45% repetitive. Despite the fragmentation, the assembly yielded high BLAST hits to 97.2% of the available EST-derived cowpea unigenes (http://harvest.ucr.edu). This number may be an underestimate of the representation of genes in IT97K-499-35 because the 17 cowpea accessions used for the EST libraries may contain genes not present in IT97K-499-35. The WGS assembly also produced BLAST hits to 24,712 common bean gene models, which is 90.9% of the total number of predicted protein-coding loci (Schmutz et al. 2014). The average GC content of the WGS assembly was 35.96%, similar to other sequenced legumes (Varshney et al. 2012; Schmutz et al. 2014; Yang et al. 2015).

**Table 2.** **WGS assembly characteristics and anchoring to the genetic map.**

| SOAPdenovo Assembly | # Scaffolds | Total Size (bp) | Non-N Size (bp) | N50 | L50 | N90 | L90 | Pv. Gene Models |
| --- | --- | --- | --- | --- | --- | --- | --- | --- |
| Scaffolds (all) | 644,126 | 754,150,495 | 323,048,341 | 6,332 | 30,516 | 420 | 148,491 | 24,712 |
| Scaffolds ($\geq 1$ kb) | 97,777 | 646,178,954 | 215,706,467 | 8,688 | 23,133 | 4,588 | 68,160 | 23,369 |
| Anchored Scaffolds (all) | 25,537 | 237,061,317 | 100,209,137 | 12,546 | 6,257 | 5,627 | 17,190 | 17,807 |
| Anchored Scaffolds ($\geq 1$ kb) | 24,342 | 236,188,142 | 99,338,135 | 12,586 | 6,223 | 5,675 | 17,051 | 17,191 |

N50: length for which the collection of scaffolds of that length or longer contains at least half of the sum of the lengths of all scaffolds in the WGS assembly.

N90: length for which the collection of scaffolds of that length or longer contains at least 90% of the sum of the lengths of all scaffolds in the WGS assembly.

L50: minimum number of scaffolds accounting for more than 50% of the WGS assembly.

L90: minimum number of scaffolds accounting for more than 90% of the WGS assembly.

Pv.: *Phaseolus vulgaris*.

### Development of the Cowpea iSelect Consortium Array

A total of 36 additional cowpea accessions relevant to Africa, China and the USA were shotgun sequenced using Illumina HiSeq 2500 (12.5x average coverage) and aligned to the WGS assembly of IT97K-499-35 to discover SNPs. These accessions were chosen to represent the geographic, phenotypic and genetic diversity of cultivated cowpea (Figure S1; Table S1). An additional set of 12.5x HiSeq data was also produced from IT97K-499-35 and included as a control against spurious SNP calls. The reads were mapped to the reference sequence (Data S1) using BWA (Li et al. 2009a) to generate a. bam file. Then, SAMtools (Li et al. 2009b), SGSautoSNP (Lorenc et al. 2012) and FreeBayes (Garrison et al. 2012) were used to generate three overlapping sets of candidate SNPs, from which the intersection yielded about 1 million SNPs. No accession contributed substantially more SNPs than any other, highlighting the broad coverage of diversity within the set of accessions used for SNP discovery. The most distant accession from IT97K-499-35 was UCR779 (differing at 25% of SNP loci) followed closely by the four Chinese accessions (22%-24%). The accessions most closely related to IT97K-499-35 were the IITA breeding lines IT89KD-288, IT93K-503-1 and IT84S-2246 (12%-13%). The set of ~1 million SNPs was filtered to 55,496 SNPs for the design of an Illumina iSelect Consortium Array (see Esperimental procedures for details on filtering criteria). The design also included 1,163 SNPs from the prior GoldenGate Assay (Muchero et al. 2009) and 60 presumed organelle SNPs, for a total of 56,719 intended SNPs (60,000 assays). From those, 51,128 SNPs (90.1%) were represented in the final product manifest (Data S2). The cowpea iSelect Consortium Array is available from Illumina (Illumina Inc., San Diego, CA; http://www.illumina.com/areas-of-interest/agrigenomics/consortia.html).

### Construction of a consensus genetic map for cowpea

Five bi-parental RIL populations were used to develop a consensus genetic map (Table S2). Monomorphic SNPs and those with an excessive number of missing and/or heterozygous calls were eliminated, as well as individuals that were duplicated or highly heterozygous. The number of lines per population used for mapping ranged from 94 to 135 (Table S2) for a total of 575 RILs. A genetic map was constructed using MSTmap (Wu et al. 2008; http://mstmap.org/) at LOD 10 for each RIL population. Linkage groups (LGs) were numbered and oriented based on a previous cowpea consensus map (Lucas et al. 2011). Individual maps and the genotype data used for their construction can be found in Data S3. Two maps (Sanzi x Vita7 and CB27 x IT82E-18) each had two chromosomes separated into two LGs (Table S2 and Data S3) due to regions where parents lack polymorphisms. One region of identity between CB27 and IT82E-18 on LG4 impacted the number of polymorphisms and marker bins, and the total size of that LG in the specific genetic map (Table 3). Genetic map sizes varied among the five populations, from 803.4 cM in ZN016 x Zhijiang282 to 917.1 cM in Sanzi x Vita7 (Table 3).

**Table 3.**
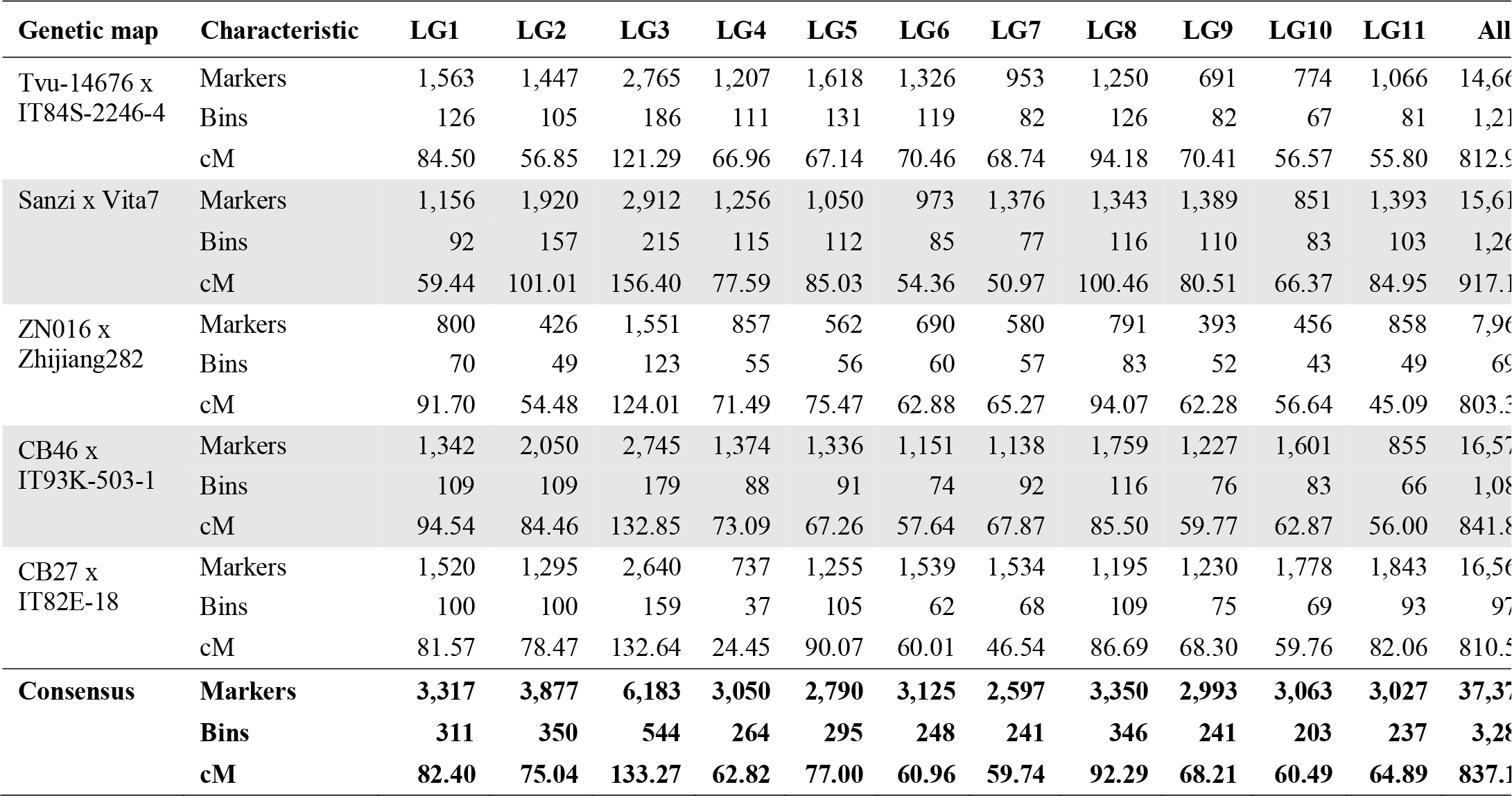
Distribution of SNPs in the individual component maps and the consensus map.

Individual maps were merged into a consensus map using MergeMap (Wu et al. 2011; http://mergemap.org/). Equal weight was given to each individual map. MergeMap identified a few conflicts in marker order, which were resolved by deleting a few conflicted markers with priority given to the map with the highest resolution in the particular LG (i.e. more bins). No SNP was placed on different LGs between maps. Since MergeMap’s coordinate calculations for a consensus map are inflated relative to cM distances in individual maps, consensus LG lengths were normalized to the mean cM length from the individual maps. The resulting consensus map contains 37,372 SNP loci mapped to 3,280 bins (Table 3; Data S4). This is a 34-fold increase in marker density and a four-fold increase in resolution (number of bins) over the consensus map of Lucas et al. (2011). The new consensus map includes 757 SNPs that were included in the prior GoldenGate assay (Muchero et al. 2009). The map spans 837.11 cM at an average density of one bin per 0.26 cM and 11.4 SNPs per bin. The new consensus map has dense coverage of all eleven cowpea LGs, with 1.85 cM on LG1 being the largest gap (Figure S2; Data S4).

### Syntenic relationships between cowpea and common bean

Similar to cowpea, common bean is a diploid member of the Phaseoleae tribe with 2n = 22 chromosomes. The iSelect SNP design sequences were compared to *P. vulgaris* gene models (Schmutz et al. 2014) to clarify the syntenic relationships of cowpea with this closely related species. The 26,550 SNPs that were mapped in *V. unguiculata* and matched a *P. vulgaris* gene model provided a view of synteny (Figure 1). Six cowpea LGs (VuLG2, VuLG6, VuLG8, VuLG9, VuLG10 and VuLG11) are largely collinear with six common bean pseudomolecules (Pv7, Pv6, Pv9, Pv11, Pv10 and Pv4, respectively), while the rest have synteny mainly with two common bean pseudomolecules (Figure 1; Table S3). From these five cowpea LGs with one-to-two relationships, three (VuLG3, VuLG4, and VuLG7) have a higher number of BLAST hits, and over a longer genome interval, with one *P. vulgaris* chromosome (Pv3, Pv1 and Pv2, respectively; Figure 1 and Table S3). The other two cowpea LGs, VuLG1 and VuLG5, both have their largest block of homologous synteny with Pv8, followed by Pv5 and Pv1, respectively (Figure 1; Table S3).

**Figure 1.**
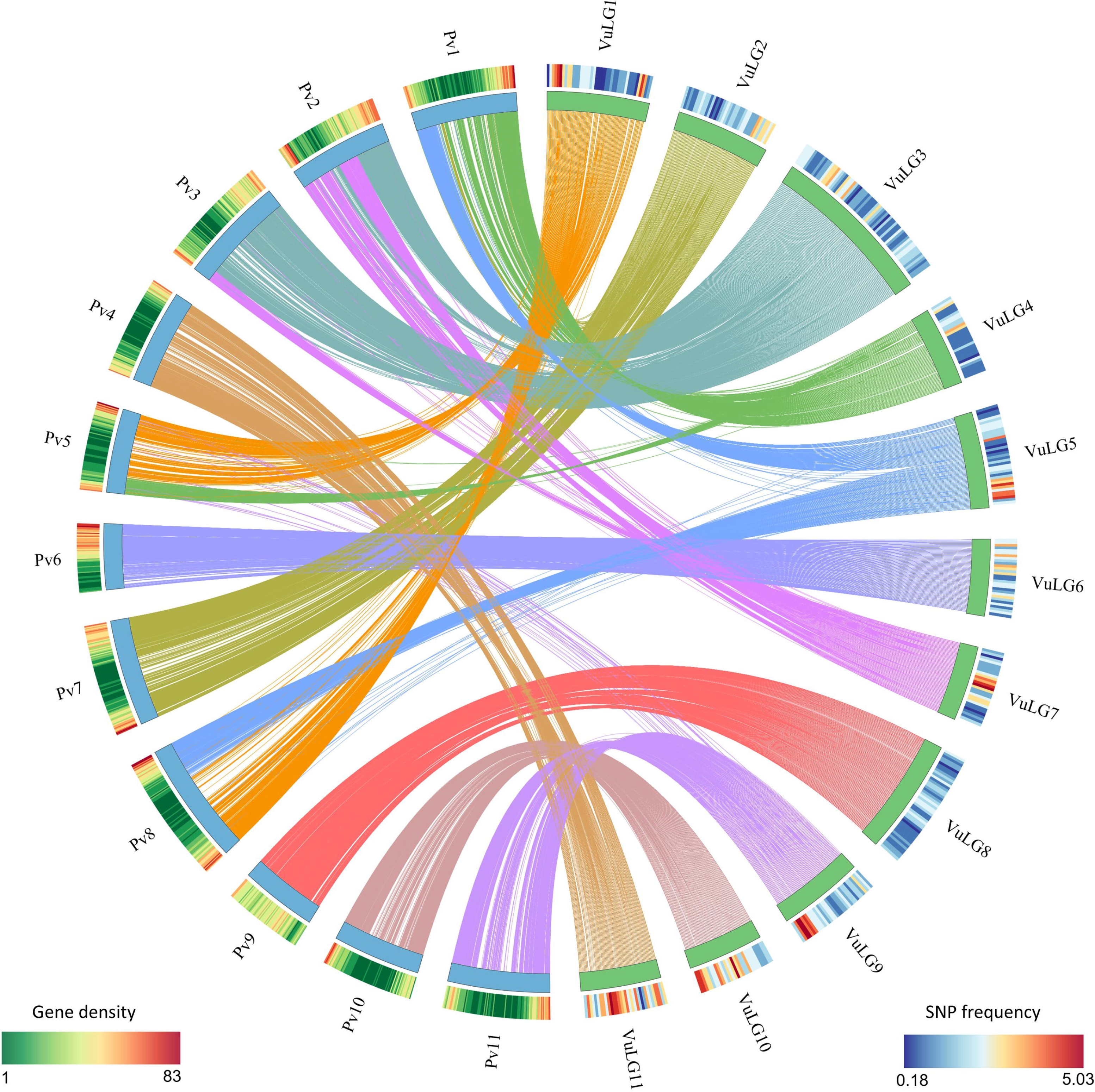
Circos illustration of synteny between cowpea linkage groups (VuLG) and common bean pseudomolecules (Pv). SNP frequencies calculated for 2 cM windows and normalized to the total anchored scaffold size (in kb) are shown for the 11 cowpea LG. *P. vulgaris* gene densities are also shown for 500 kb windows.

The same numbering scheme for common bean and cowpea chromosomes would facilitate comparative studies between the two species. Adoption of the chromosome numbers of *P. vulgaris* according to synteny relationships with LGs of cowpea seems sensible, but additional cowpea sequence information will be needed to clarify the relationships between VuLG1 and VuLG5 with Pv1, Pv5 and Pv8. The BAC-FISH analysis by Iwata-Otsubo et al. (2016) that correlates the genetic and chromosome maps in cowpea can be used to orient the cowpea genetic map so that it meets the convention of displaying the short arm on top.

### Genetic anchoring of BACs and WGS scaffolds

The 37,372-SNP consensus map was used to anchor WGS and BAC assemblies to genetic map positions. The iSelect SNP design sequences were used as BLAST queries to search against WGS and BAC sequences, and matches with an e-value = 1e^−50^ or better were tallied. Assembled sequences were considered anchored to the genetic map if 100% of the matching SNPs mapped to the same LG, and were at most 5 cM apart (Data S5 and S6). The anchored sequences contain 420 Mb of BAC assemblies (Table 1 and Data S5) and 100 Mb of the WGS assembly (237 Mb scaffold size including Ns; Table 2 and Data S6). For BACs, this is an overestimate of the actual genome coverage because BAC sequences have ~23% overlap (see “Physical map and BAC sequencing”), resulting in a reduced estimate of 323 Mb of unique sequences within anchored BACs. Also, observe in Table 2 that 95.3% of the anchored WGS scaffolds are larger than 1 kb and they comprise 99.1% of the anchored non-N sequence. Thus, the anchored portion of the WGS assembly, which is comprised mainly of 24,342 scaffolds larger than 1 kb among 25,537 anchored scaffolds, contains many fewer fragments than the entire WGS assembly (644,126 scaffolds). This is the outcome of having selected SNPs in the largest WGS contigs as a final criterion in SNP selection (see Experimental procedures).

### Distribution of genetic variation

The anchoring of WGS scaffolds to the genetic map enabled investigation of the frequency and positional distribution of genetic diversity in the cowpea genetic map. Nearly half of the 1,036,981 SNPs discovered from the 36 diverse cowpea accessions were anchored to the genetic map based on the anchoring of 25,537 WGS scaffolds using mapped iSelect SNPs. This information was used to examine the SNP frequency and distribution across the eleven cowpea LGs. Frequencies were calculated for 2 cM intervals and normalized to the total anchored scaffold size. SNP frequencies were not uniformly distributed across the genetic map (Figure 1). LG11 and LG10 had significantly higher SNP frequencies than all other cowpea linkage groups. Relatively higher SNP frequencies were also observed in the distal ends of LG5 and LG9, in the centromeric region of LG7, and toward the ends of LG1. In contrast, LG8 had relatively low SNP diversity (Figure 1). There was no clear relationship between the most diverse cowpea genomic regions and the gene-dense syntenic regions of common bean (Figure 1).

### Genetic diversity in West African breeding programs

Evaluation of the extent and distribution of genetic diversity has important implications for breeding programs and for the conservation of genetic resources. The level of genetic diversity within West African cowpea breeding programs was examined by comparing the diversity among cultivars/breeding lines with the diversity among locally adapted landraces. A total of 146 unique cultivated cowpea accessions were genotyped, including 105 cultivars or breeding lines from the breeding programs of IITA (Nigeria), INERA (Institut de l'Environnement et de Recherches Agricoles, Burkina Faso), ISRA (Institut Senegalais de Recherches Agricoles, Senegal), and CSIR-SARI (Council for Scientific and Industrial Research, Savanna Agricultural Research Institute, Ghana), and 41 landraces collected from these same countries (Table S4). A total of 46,620 SNPs were polymorphic among these accessions and were used for the diversity analysis.

Principal Component Analysis (PCA) was performed to clarify the genetic relationships between accessions. PCA clearly shows that cultivars and breeding lines do not cluster by breeding program, but rather are scattered in the PC plot (Figure 2). This indicates that the four West African breeding programs are working with very similar materials, except for somewhat narrower diversity within the Ghana program than in the other three West African programs. Also, landraces were not differentiated from the improved breeding materials, indicating limited impact of breeding on the overall allelic composition. Landraces were less dispersed than cultivars/breeding lines, mostly distributed along the first PC. The landraces from Senegal were the most diverse within the landrace group. Thirteen landraces that were collected in the same geographical area of Burkina Faso clustered together (Figure 2), indicating high genetic similarity between them. Ghanaian and Nigerian landraces have sparse representation in this analysis.

**Figure 2.**
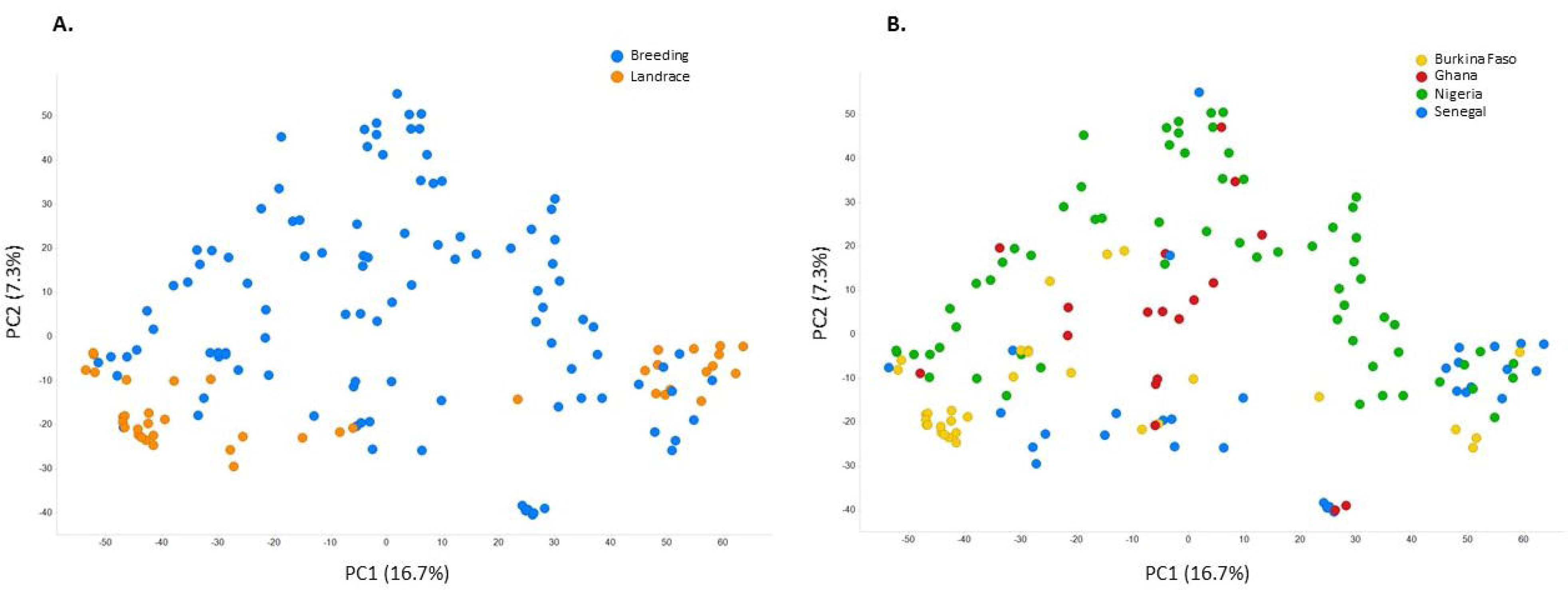
Principal Component Analysis (PCA) of 146 cultivated cowpea accessions from West Africa. The plot on the left (A) displays accessions colored by the type of material (breeding vs. landrace), while the plot on the right (B) show accessions colored by their country of origin.

Polymorphism information content (PIC; Botstein et al. 1980) was used as an evaluation of genetic diversity within the 146 West African accessions, and between breeding materials and landraces. PIC values ranged from 0.007 to 0.375 (0.375 is the maximum PIC for bialellic markers), with an average of 0.247 when considering all 146 accessions. The average PIC value among cultivars and breeding lines was slightly higher than that in landraces (PIC = 0.242 and 0.234, respectively). However, given the small sample size of local landraces, this may be an inaccurate estimate of the diversity in West African landrace germplasm. The spatial patterns of genetic diversity across the eleven LGs were explored by plotting PIC values for landraces and improved breeding lines against the consensus genetic map (Figure 3; Figure S3). Regions of very low average PIC values (<0.05) were identified for linkage groups 1, 3, 4, 7 and 8. Many alleles in those regions were fixed (PIC = 0) in breeding materials, landraces, or both (Table 4; Data S7). Three of those regions coincide with the positions of loci for traits involved in subspecies differentiation [pod length (Xu et al. 2016), LG1 and LG4; Table 4] or environmental adaptation [heat tolerance (Lucas et al. 2013a), LG3; Table 4], whereas traits associated with other low PIC regions are unknown (Figure 3; Figure S3; Table 4). A moderate reduction of diversity was observed in a region on LG5 where a domestication-related QTL, *Css-1*, involved in seed size (Lucas et al. 2013b) has been mapped (Figure S3).

**Figure 3.**
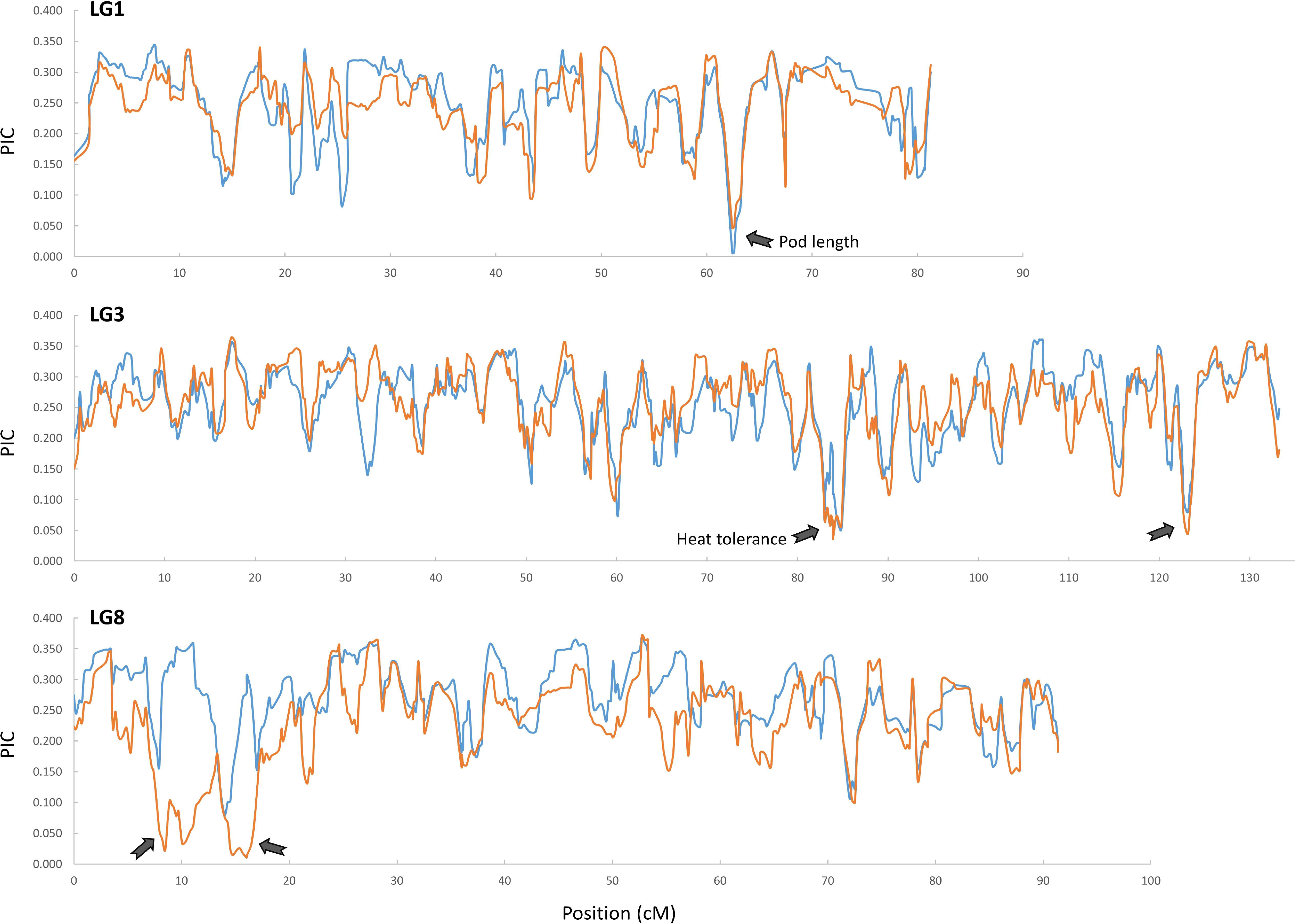
Diversity in West African cultivated cowpeas across linkage groups 1, 3, and 8. Blue lines represent polymorphism information content (PIC) values calculated in cultivars and breeding lines, while orange lines represent PIC values in local landraces from Burkina Faso, Ghana, Nigeria and Senegal. PIC was averaged across a sliding window of 5 genetic bins with a step of one bin. Arrows indicate regions with a markedly depletion of genetic diversity in one or both types of materials.

**Table 4.** Regions showing the lowest genetic diversity in West African improved breeding lines, local landraces, or both. Minimum Polymorphism Information Content (PIC) values per bin are shown for the breeding and landrace group, as well as the number of SNPs contained in those regions and any QTL previously identified in those regions.

| LG | cM range | Avg. Min PIC<br>Breeding | Avg. Min PIC<br>Landraces | # SNPs in<br>range | QTL |
| --- | --- | --- | --- | --- | --- |
| 1 | 61.83 - 63.80 | 0.000 | 0.038 | 97 | Pod Length (Xu et al. 2016) |
| 3 | 82.97 - 85.69 | 0.000 | 0.000 | 178 | Heat tolerance (Lucas et al. 2013a) |
| 3 | 122.73 - 124.33 | 0.019 | 0.000 | 49 | — |
| 4 | 7.40 - 8.83 | 0.000 | 0.046 | 490 | Pod Length (Xu et al. 2016) |
| 7 | 50.53 – 51.17 | 0.028 | 0.000 | 13 | — |
| 8 | 8.46 - 11.95 | 0.262 | 0.000 | 67 | — |
| 8 | 15.69 - 17.43 | 0.121 | 0.000 | 39 | — |

Patterns of polymorphism in breeding materials were similar to those in landraces throughout most of the genetic map. This is consistent with their close genetic relationships illustrated by the PCA analysis (Figure 2). However, there were some notable differences. The most distinct difference was in LG8, where West African landraces showed very low levels of genetic diversity while improved breeding materials exhibited very high PIC values (Figure 3), suggesting that additional diversity may have been introduced into the breeding programs for one or more traits in this region.

## DISCUSSION

Increase in climate variability is projected to have the greatest negative consequences on agricultural and human systems in the tropical and subtropical developing world, aggravating food insecurity in already vulnerable populations (Thornton et al. 2014). Cowpea is a relatively drought and heat tolerant crop that provides protein to nearly 200 million Africans and cash income to smallholder farmers (Thomson et al. 2008). The limited availability of genome resources for cowpea has contributed to the relatively slow development of higher yielding varieties adapted to tolerate abiotic and biotic stresses. This report presents 323 Mb of WGS and 497 Mb of BAC sequence information, a tool to simultaneously test 51,128 single nucleotide variants, and a high-density genetic map providing coordinates for most of those sequences and variants. Application of these resources can be made for genome-wide association studies (GWAS) of cowpea germplasm to discover favorable alleles for simple and complex traits, as already being conducted in other legume crops (e.g. Kujur et al. 2015; Ray et al. 2015). Useful variation can then be connected to assembled genome sequences-including BACs-annotated for *P. vulgaris* syntenic gene models, thereby increasing the precision and speed of cowpea improvement.

One of the biggest obstacles in comparing and using results obtained by different research groups is the lack of a common nomenclature for cowpea linkage groups. With a high SNP coverage of the genome and connections to cowpea genome sequences, this study provides the basis for a unified chromosome nomenclature for the cowpea research community. Such common nomenclature could adopt the *P. vulgaris* chromosome numbering on the basis of synteny comparisons between both species as well as cytogenetic studies in cowpea (Iwata-Otsubo et al. 2016) and between cowpea and common bean (Vasconcelos et al. 2015). While several cowpea LGs are largely syntenic with one *P. vulgaris* chromosome, further resolution is needed to satisfy a single nomenclature for those LGs whose syntenic relationships with common bean are less clear. The goal would be to extend a standard chromosome numbering to other diploid *Vigna* species whose genomes have been sequenced and are integrated into a genome database (Sakai et al. 2016). This would facilitate the transfer of genomic information on target traits from one Fabaceae species to another.

West Africa is the region with the largest production and consumption of cowpea in the world (FAOSTAT 2012, Singh 2014). Evaluating the wealth of genetic diversity present in the West African breeding germplasm is important to manage breeding programs and assure future genetic gains. By applying the Cowpea iSelect Consortium Array to 146 breeding lines and landraces, we provided a useful overview of genetic variability in West African cultivated germplasm. We found moderately high levels of genetic diversity in breeding materials (PIC ~ 0.25), which are similar to those present in regional landraces. This indicates that, at the genome scale, the 40+ years of breeding practices in the region did not deplete diversity available in the West African breeding germplasm. Introgression of Asian and US germplasm into West African breeding lines was reported previously (Fang et al. 2007). This would explain the relatively high levels of diversity in breeding materials and their more extended distribution along PC2 (Figure 2). The common agro-ecological zones which extend across cowpea production areas of the four included countries of Burkina Faso, Ghana, Nigeria and Senegal facilitates exchange of germplasm between breeding programs and supports their genetic relatedness. The influence of the IITA, Nigeria breeding program, in being a regional distributor of new breeding materials during the last few decades is difficult to quantify, but this has no doubt contributed significantly to the overall similarity of the breeding materials across these West African national breeding programs. Even though the sample size is relatively small, the lack of strong population structure in this West African dataset would be beneficial for identifying true marker-trait associations via GWAS, provided that phenotypic diversity is present for the traits of interest (Hamblin et al. 2011; Korte and Farlow 2013).

Average PIC values should be interpreted cautiously because patterns of polymorphism vary across LGs. In fact, although the overall genetic diversity within the West African breeding population is relatively high, we identified genomic regions of diversity depletion. Those regions may contain favorable alleles for important traits that became fixed during domestication and breeding selection. The lowest PIC values in LG1 coincide with the position of SNPs associated with pod length in Chinese germplasm of *V. unguiculata* subspecies *sesquipedalis* (Xu et al. 2016). One interpretation could be that there has been selection for shorter-podded forms in West Africa, a trait which correlates with bushy plant architecture and increased pod number per plant in cultivated dry grain cowpeas (Bapna et al. 1972). Also, a previously reported QTL for heat tolerance *(Cht-5*) coincides with a very low-diversity region of LG3 (Lucas et al. 2013a). Favorable alleles at this QTL were donated by the line IT82E-18, the African parent of the RIL population (Lucas et al. 2013a; Table 3). The low diversity in this region of LG3 may reflect selection at this genomic region for better yield performance of West African cowpeas under higher growing season temperatures. A more complete analysis of diversity in cultivated and wild cowpea germplasm is required to further test these hypotheses of trait association with genome regions of low genetic diversity, and to reveal if other low-diversity regions contain traits involved in domestication or crop improvement.

The release of PIC values for each SNP (Supplementary Table 9) is another valuable resource stemming from this work. PIC values can be used as criteria for selecting efficient subsets of markers for conversion to other platforms. Customized, maximally informative subsets of markers have numerous applications including routine tests of seed purity, validation of germplasm fidelity, verification of successful crosses and guidance of progeny selection in later generations during trait introgression into preferred backgrounds via backcrossing.

The new cowpea genome resources must be easily accessed if they are to be widely utilized for basic research and agricultural development. All information presented in this manuscript, including SNPs, BAC and WGS sequences, *Phaseolus vulgaris* gene annotations, as well as genetic anchoring information are available through the online interface HarvEST:Web (http://harvest-web.org/hweb/utilmenu.wc) or in the latest version of HarvEST:Cowpea (http://harvest.ucr.edu). In addition, a cowpea-common bean synteny viewer has been implemented in HarvEST:Cowpea, enabling facile comparisons between these two closely related legume crops.

## EXPERIMENTAL PROCEDURES

### Physical mapping and BAC-end sequencing

Cowpea accession IT97K-499-35 was grown for three generations by single seed descent and then increased to provide a supply of seed for DNA isolation. The material was screened with the Illumina GoldenGate assgay (Muchero et al. 2009) to establish that homozygosity was attained. Young seedling leaves were harvested at UCR and shipped on dry ice to Amplicon Express (Pullman, WA) for purification of nuclei and extraction of mainly nuclear DNA. Two BAC libraries were then constructed by Amplicon Express (Pullman, WA) from high molecular weight DNA of cowpea accession IT97K-499-35 using restriction enzymes *HindIII* and *MboI*. After partial digestion with restriction enzymes, high MW cowpea DNA fragments were ligated with *HindIII* or *BamHI* linearized BAC vector pCC1. Ligated DNA molecules were introduced into *Escherichia coli* DH10B cells by electroporation and plated on LB agar containing 12.5 mg/ml chloramphenicol, 0.5 mM IPTG and 40 mg/ml X-Gal and cultured overnight. White colonies were picked and inoculated into 384-well plates containing LB freezing buffer. Cultures were incubated at 37°C for 24 h with aerations, and then stored at-80°C.The libraries contained 36,864 clones each, with average insert sizes of 150 kb for the *HindIII* library and 130 kb for the *Mbo*I library.

BAC clones from the two libraries (36,096 from *HindIII* and 23,312 from *MboI*) were fingerprinted using the SNaPshot-based fingerprinting procedure (Luo et al. 2003). BAC DNAs were simultaneously digested with five restriction enzymes (BamH1, EcoRI, *XbaI, XhoI*, and HaeIII), and then labeled with the SNaPshot labeling kit (Luo et al. 2003). The fragments were sized on an ABI3730XL instrument with the GS1200Liz size-standard (Gu et al. 2009). Fragment sizes in the range of 100-1000 bp were compiled for computational assembly. After removing substandard fingerprints, potential cross contamination and clones with less than 40 total fragments, fingerprints from 43,717 clones (73.6%) were used for an initial contig assembly using the FPC software (Soderlund et al. 2000). This initial assembly was performed with a relatively high stringency (1×10^−45^) to minimize co-assembly of clones from unrelated regions of the genome. The “DQer” function of the FPC software was used for second stage assembly by disassembling contigs containing more than 15% questionable clones. The “Single-to-End” and “End-to-End” merging function of FPC was used for a final, third stage assembly by stepwise decreases of assembly stringency based on Sulston score cutoff values (down to 1×10^−35^). Finally, the 10% largest contigs were subjected to manual editing, examining with CB map analysis and disjoining contigs with CB analysis results at 1×10^−30^.

The same BAC DNA used for fingerprinting was also used for BES. BAC clones were sequenced using pIndigoBAC5 Reverse End-Sequencing primer (5′ TACGCCAAGCTATTTAGGTGAGA 3′) and BigDye terminator chemistry (Applied Biosystems) on an ABI3730XL automated sequencer (Applied Biosystems). Raw sequence reads were trimmed with Phred program using a quality score of 20 (Ewing and Green, 1998). BES from vector sequences, E *coli*, mitochondria and chloroplasts were identified using BLASTN. The chloroplast sequences of common bean (DQ886273.1), soybean (DQ317523), *Medicago truncatula* (AC093544), *Lotus japonicus* (AP002983), and mitochondrial DNA sequences of Arabidopsis (Y08501.2) and rice (DQ167399.1) were used to identify organelle contaminations. The resulting high quality BES were then processed with the RepeatMasker program (www.repeatmasker.org) to identify characterized repeats. Cowpea BES with more than 80% of the sequence length showing homology to known repeats were removed, otherwise the BES were kept but the repetitive region was marked using letter N. Self-comparisons were conducted with the RepeatMasker processed sequences to further filter the cowpea-specific repeat elements.

### MTP sequencing and BAC assembly

A set of minimal tiling path BACs was chosen using the FMTP method of Bozdag et al. (2013). MTP BACs were paired-end sequenced (2 × 100 bases) using Illumina HiSeq2000 (Illumina, Inc, San Diego, CA, USA). Sequencing was done in two sets of 2,197 BACs (Vu1 and Vu2) applying a combinatorial pooling design (Lonardi et al. 2013). After quality-trimming, reads in each pool were “sliced” into smaller samples of optimal size, deconvoluted, and then assembled BAC-by-BAC using SPAdes (Bankevich et al. 2012), as explained in detailed by Lonardi et al. (2015). From the 4,394 intended BACs, 4,355 produced sufficient reads to generate an assembly. Raw reads for these cowpea BACs were deposited in NCBI SRA under accession numbers SRA052227 and SRA052228.

To estimate the percentage of overlapping BAC sequences, 19-mers occurring at least 4 times were identified and used for repeat-masking of sequences. Repeat-masked sequences were then BLASTed against themselves using an e-value cutoff of e^−40^. Only overlapping sequences > 300 bp were considered to be overlaps. To estimate the gene content of the BAC assemblies, BAC sequences were compared to cowpea EST-derived “unigenes” (http://harvest.ucr.edu) and *Phaseolus vulgaris* gene models (Schmutz et al. 2014) using BLAST (e-value cutoffs of e^−40^ and e^−25^, respectively).

### Whole-genome shotgun sequencing and assembly

The same batch of IT97K-499-35 nuclear DNA that was used for BAC library construction was used for whole-genome shotgun sequencing. About 394 M paired-end reads (equivalent to ~65x coverage) with an average read length of ~100 bases after quality-trimming were produced at the National Center for Genome Resources (NCGR; Santa Fe, NM) on an Illumina GAII sequencing instrument. An additional ~90 M Illumina reads were produced using an Illumina Hi-Seq sequencing instrument at NCGR from one 5 kb long-insert paired end (LIPE) library made from the same batch of nuclear DNA.

For the assembly, two additional sets of Sanger sequences were included. One set of Sanger sequences were the basis of a prior publication on “gene-space sequences” (GSS; Timko et al. 2008), comprised of ~250,000 reads from methyl filtered fragments of IT97K-499-35. The other set of Sanger sequences included the BAC-end sequences described above. The assembly combined the paired-end short reads, LIPE, GSS, and BES data using SOAPdenovo with k=31 (Luo et al. 2012). To estimate the gene content of the WGS assembly, sequences were BLASTed against cowpea EST-derived “unigenes” (http://harvest.ucr.edu) and *Phaseolus vulgaris* gene models (Schmutz et al. 2014), using e-value cutoffs of e^−40^ and e^−25^, respectively.

### SNP discovery and design of the Cowpea iSelect Consortium Array

A total of 32 accessions were sequenced to 12.5x coverage by the Beijing Genomics Institute (BGI) using Illumina HiSeq 2500 (Illumina, Inc., California, USA). Four additional accessions from China (see Table S1) were sequenced at the Majorbio Pharm Technology Co. Ltd (Shanghai). Additional sequences of IT97K-499-35 were produced in the Genomics Core Facility at the University of California, Riverside.

The WGS assembly from IT97K-499-35 described above was used as the reference to map each of these 36 sets of reads, and one new set of HiSeq sequences from the reference genotype sequenced at the University of California Riverside (Genomics Core Facility). This 37^th^ set was used as a control (i.e. SNPs call in this accession were considered false positives). BWA (Li et al. 2009a) was used to uniquely map each set of reads (BWA mem with-M option to mark shorter split hits as secondary). Reads which mapped to multiple locations were excluded from further analysis. Alignment files were merged with the software tool Picard to a single “sam” file. Reads that “hanged off” the end of the contigs in the reference sequence were clipped with Picard. Also, to avoid skewed variant calling, duplicated reads were marked with Picard.

To filter putative SNPs to a shorter list of highest confidence variants, three software packages were used, namely SAMtools (Li et al. 2009b), SGSautoSNP (Lorenc et al. 2012), and FreeBayes (Garrison et al. 2012). It was not possible to utilize GATK (McKenna et al. 2010) because it requires a relatively large set of confirmed training SNPs for the base quality score recalibration phase, and no such set of SNPs was available for cowpea. In total, SAMtools discovered 5,108,787 SNPs using mpileup with default parameters, SGSautoSNP detected 2,488,797 SNPs and FreeBayes called a total of 8,269,140 SNPs. Finally, we used the haplotype-based variant detection tool FreeBayes to independently call the cowpea SNPs. FreeBayes called a total of 8,269,140 SNPs on the cowpea genome. An intersection set of SNPs was then identified, leading to 1,036,981 SNPs that were identified by all three methods. Additional filtering was required to reduce the number of SNPs to the target density of 60,000 SNP assays designed as a community resource for future germplasm characterization. These filtering steps included: 1) designability score based upon Illumina’s Assay Design Tool; 2) avoidance of a SNP whose adjacent sequences occurred frequently in the genome assembly; 3) consideration of allele frequency, generally avoiding SNPs with only one accession carrying the minor allele; 4) selection of two SNPs in or near each inferred cowpea gene based on MUMmer sequence alignment with *Phaseolus vulgaris* gene models (Schmutz et al. 2014); 5) requirement for a minimum distance from a SNP that had already been selected, 6) preference against an A/T or C/G SNP since these require two beadtypes (assay space), and 7) location within a relatively larger WGS contig to maximize the amount of WGS contigs that could subsequently be anchored to a SNP-based genetic map.

In addition to SNPs discovered by WGS sequencing of diverse accessions, a total of 1,163 SNPs previously validated on the GoldenGate platform (Muchero et al. 2009) were included in the design to facilitate comparisons with prior genotyping research. A total of 56,719 SNPs were submitted for assay design using 60,000 beadtypes with 51,128 assays included in the final manifest for the publicly available Cowpea iSelect Consortium Array.

### Consensus genetic map construction

Five bi-parental RIL populations developed previously (Muchero et al. 2009; Lucas et al. 2011) were genotyped with the Cowpea iSelect Consortium Array at the University of Southern California. SNPs were called using the GenomeStudio software (Illumina, Inc.). To meet assumptions of the clustering algorithm, “synthetic heterozygotes” were constructed and included in the initial set of 96 genotyped samples by creating 1:1 mixtures of DNA samples from individuals known from prior work to be most genetically distant from each other. The data from these individuals provided the signal needed for the algorithm to place a cluster position for heterozygotes. SNPs with low GenTrain scores were visually inspected based upon manufacturers published best practice for optimizing accuracy in genotyping projects (http://www.illumina.com/documents/products/technotes/technote_infinium_genotyping_data_analysis.pdf). The resulting cluster file is available upon request.

SNP data from each population were exported from GenomeStudio and curated to eliminate 1) monomorphic SNP loci, 2) SNPs with >20% missing or heterozygous calls, and 3) segregation-distorted markers (MAF<0.25). RILs were also curated to remove individuals with >10% heterozygous loci or those carrying many non-parental alleles. Identical individuals were also thinned to one such individual prior to mapping. Genetic maps for each RIL population were constructed at LOD 10 using MSTmap (Wu et al. 2008; http://mstmap.org/). Because the level of residual heterozygosity varied among populations, different population type options were chosen for map construction in MSTmap (RIL 7 for Tvu-14676 x IT84S-2246-4; RIL 6 for Sanzi x Vita7 and ZN016 x Zhijiang282; and RIL5 for CB46 x IT93K-503-1 and CB27 x IT82E-18). Other parameters for MSTmap included: grouping LOD criteria = 10; no mapping size threshold = 2; no mapping distance threshold = 10 cM; try to detect genotyping errors = no; and genetic mapping function = kosambi. Output maps were inspected to identify and remove data that would result in presumably spurious double recombination events, unless supported by several markers or moderate to large genetic distances.

LGs from each population were numbered and oriented based on the previous cowpea consensus map (Lucas et al. 2011) and then merged into a consensus map using MergeMap (Wu et al. 2011; http://mergemap.org/). Equal weight was given to each individual map (weight = 1.0). MergeMap identified a few conflicts in marker order, which were resolved by deleting a few conflicted markers with priority given to the map with the highest resolution in the particular LG (i.e. more bins). Since MergeMap’s coordinate calculations for a consensus map are inflated relative to cM distances in individual maps, consensus LG lengths were normalized to the mean cM length from the individual maps.

### Synteny with *Phaseolus vulgaris*

The cowpea genome assembly described above was compared to *P. vulgaris* pseudomolecules and unanchored scaffolds (from https://phytozome.jgi.doe.gov/pz/portal.html) using MUMmer (Kurtz et al. 2004). Alignments that were further used had a minimum identity of 55.11% and a mean identity of 89.24%. The positions of *P. vulgaris* gene models within the aligned regions was used to position each cowpea SNP relative to *P. vulgaris* gene models. A synteny plot was constructed based on SNPs that had a cM position in the cowpea consensus map and fell within the region of the cowpea sequence that was aligned with a common bean gene model. Circos v.67-7 (Krzywinsk et al. 2009) was used to illustrate the synteny between each cowpea linkage group and common bean chromosome that shared 50 or more SNPs. Cowpea LGs were plotted according to cM lengths, while common bean chromosomes were plotted as physical length.

Cowpea SNP frequencies were based on the number of discovered SNPs per genetic bin and the total size of the WGS scaffolds allocated into the corresponding bin. Then, for every 2 cM window the number of SNPs allocated within that window was divided by the sum of the corresponding WGS scaffold sizes in kb. Two outlying values were replacing by a maximum value so that all of the other calculated values could be easily visualized. *Phaseolus vulgaris* gene densities were calculated as number of genes available from Schmutz et al. (2014) per 500 kb windows.

### Genetic analysis of West African accessions

A total of 146 accessions were genotyped with the Cowpea iSelect Consortium Array. Monomorphic loci were eliminated, as were SNPs with missing or heterozygous calls in more than 20% of the samples. PCA was conducted in TASSEL v5.0 (Bradbury et al. 2007; http://www.maizegenetics.net) using SNPs with MAF>0.05, and results were displayed using TIBCO Spotfire^®^ 6.5.0. PIC values were calculated using the method of Botstein et al. (1980) for all 46,620 SNPs in the entire set of samples, and then separately in the cultivar/breeding lines and landraces. PIC values were plotted along the consensus genetic map by averaging PIC values across a sliding window of 5 bins in 1 bin steps.

## ACCESSION NUMBERS

The BAC sequence reads supporting the results of this article are available in NCBI SRA under accession numbers SRA052227 and SRA052228.

## ACKNOWLEDGMENTS

Authors thank John Weger (Genome Core Facility, UC Riverside) and Greg D. May (NCGR) for sequencing services, and Sassoum Lo and Savanah St. Clair (UC Riverside) for DNA isolation. Development of the cowpea iSelect was supported by the Feed the Future Innovation Lab for Climate Resilient Cowpea (USAID Cooperative Agreement AID-OAA-A-13-00070) and the 2014 Illumina Agricultural Greater Good Initiative. BAC library construction, physical mapping and BAC-end sequencing was supported by the Generation Challenge Program “Tropical Legumes 1” project. Minimal tiling path BAC sequencing was supported by NSF IIS-1526742 (“III: Small: Algorithms for Genome Assembly”). Partial support was also provided by Hatch Project CA-R-BPS-5306-H. “GSS” sequencing (Timko et al. 2008) was supported by Kirkhouse Trust.

## SUPPORTING INFORMATION

**Figure S1.** Principal component analysis (PCA) of 729 samples representing the diversity of cultivated cowpea (in blue) and distribution of 34 out of the 36 accessions included in the SNP discovery panel (in red). The reference genome IT97K-499-35 is represented in green. PCA was performed using previous information from 1,535 GoldenGate SNPs.

**Figure S2.** Graphical representation of the iSelect SNP consensus genetic map for cowpea. Each horizontal line is a bin.

**Figure S3.** Polymorphism information content (PIC) values along linkage groups 2, 4, 5, 6, 7, 9, 10 and 11. Blue lines PIC values calculated in cultivars/breeding lines from four breeding programs, while orange lines represent PIC values in local landraces from Burkina Faso, Ghana, Nigeria and Senegal. PIC was averaged across a sliding window of 5 genetic bins with a step of one bin. Wide arrows indicate regions with a markedly depletion of genetic diversity in one or both types of materials, while narrow arrows represents reduced diversity regions containing a known QTL.

**Table S1.** Information of the cowpea accessions used for SNP discovery. The reference genome is underlined. UCR = University of California Riverside.

**Table S2.** Information on the individual mapping population data used for consensus map construction.

**Table S3.** Pairwise counts of the number of links between cowpea linkage groups (VuLG) and common bean pseudomolecules (Pv). Clear relationships between VuLGs and Pvs are marked in bold, while conflicting relationships are marked in red.

**Table S4.** Information of West African accession used in the study. IITA = the International Institute of Tropical Agriculture; INERA = Institut de l'Environnement et de Recherches Agricoles; ISRA = Institut Senegalais de Recherches Agricoles; SARI = the Savanna Agricultural Research Institute.

**Data S1.** Mapping statistics for 37 cowpea accessions.

**Data S2.** Information of SNPs included in the final Cowpea iSelect Consortium Assay. Pv = *Phaseolus vulgaris*, Ath = *Arabidopsis thaliana*.

**Data S3.** Five individual genetic maps (each sheet) and the genotype dataset used for their construction. “A” and “B” represent Parent 1 and Parent 2 in the cross, respectively. Lower case letters mean that the genotype calls were reversed based on the parental alleles.

**Data S4.** iSelect SNP consensus genetic map for cowpea.

**Data S5.** List of sequenced BACs and their genetic anchoring information.

**Data S6.** List of WGS scaffolds and their genetic anchoring information.

**Data S7.** Polymorphism Information Content (PIC) values considering the entire sample set (146 West African accessions; PICall), only cultivar/breeding lines (105 accessions; PICbreeding), and only landraces (41 accessions; PIClandraces).

